# Discovery of a novel flagellar filament system underpinning *Leishmania* adhesion to surfaces

**DOI:** 10.1101/2025.03.03.641248

**Authors:** Barrack O Owino, Ryuji Yanase, Alan O. Marron, Flavia Moreira-Leite, Sue Vaughan, Jack D Sunter

## Abstract

Adhesion to surfaces is a common strategy employed across biology, especially by pathogens. Within their sand fly vector, *Leishmania* parasites undergo multiple developmental stages, including the understudied haptomonad form, which adheres to the sand fly stomodeal valve via a highly modified flagellum. This adhesion, likely critical for efficient transmission, is mediated by a complex adhesion plaque from which filaments in the modified flagellum extend towards the cell body and likely connect to the flagellum attachment zone (FAZ), a cytoskeletal structure important for cell morphogenesis. However, the role of the FAZ in adhesion and its relationship with the Kinetoplastid-Insect Adhesion Proteins (KIAPs) and the filamentous structures of the plaque itself remains unclear. Here, to examine the role of the FAZ in adhesion, we generated FAZ2, FAZ5, and FAZ34 deletion mutants. Deletion of any of these FAZ proteins impaired parasite adhesion *in vitro*. Furthermore, we identified a novel and distinct set of extra-axonemal flagellar filaments important for adhesion and demonstrated that KIAP2 is an essential component of these filaments. Our findings underscore the importance of a robust connection from the cell body to the adhesion plaque for stable *Leishmania* adhesion via the highly modified flagellum.

## Introduction

A common approach of pathogens to avoid clearance and promote transmission is to anchor themselves to tissues and surfaces. *Yersinia pestis*, the plague bacterium, is a well-studied example, as it proliferates in the gut of the flea as an attached biofilm [1]. The unicellular eukaryotic parasite, *Leishmania* also adopts this strategy in its insect vector. *Leishmania* spp. are unicellular protozoan parasites transmitted by an infected sand fly and are of public health importance as the causative agents of leishmaniasis.

Leishmaniasis has a broad geographic distribution, affecting approximately 700,000 to 1 million people worldwide annually [2], with a range of clinical symptoms from disfiguring skin lesions to severe infections of the internal body organs [3–5]. There are few drugs to treat leishmaniasis, most of which have severe side effects and overreliance on them has led to a global increase in drug resistance [6].

*Leishmania* has a complex life cycle, with distinct morphological forms as it cycles between a vertebrate host and a sand fly vector [7–10]. In the sand fly, the parasite undergoes multiple developmental stages, including two adhered forms, with the nectomonad form transiently adhering to the microvilli of the midgut and the haptomonad form stably adhering to the cuticular lining of the stomodeal valve [9,11–14]. Haptomonad adhesion to the stomodeal valve contributes to *Leishmania* transmission through two interconnected phenomena. Firstly, haptomonads contribute to promastigote secretory gel production that, along with the damage adhesion causes to the stomodeal valve, results in more frequent but incomplete sand fly feeding [8,13,15–17]. Secondly, haptomonads can undergo asymmetric division to generate free swimming forms, contributing to persistent infections in the sand fly [18]. Moreover, recent work has shown that detached haptomonads are infective in their own right [14].

The promastigote morphology of the free swimming *Leishmania* parasite is characterised by the positioning of the concatenated mitochondrial DNA (kinetoplast) to the anterior of the nucleus with a long motile flagellum emerging from the anterior cell tip. The flagellum is assembled from the basal body which is physically connected to the kinetoplast. There is an invagination of the cell membrane around the base of the flagellum called the flagellar pocket, which is made up of a bulbous domain proximal to the kinetoplast and a flagellar pocket neck domain, extending to the anterior cell tip in which the membrane of the cell body is close to the flagellum. Along one side of the flagellar pocket neck, the flagellum cytoskeleton is laterally connected to the cell body cytoskeleton through the cell body and flagellum membranes by a complex cytoskeletal structure called the Flagellum Attachment Zone (FAZ) [19,20].

The FAZ is organised into three major domains (cell body, intermembrane, and flagellum domains) and FAZ proteins that localise to each domain have been identified [21,22].

Studies of these proteins have shed light on the functional significance of the FAZ in *Leishmania* biology [19,20,22,23]. FAZ5, a key component of the intermembrane domain, was shown to be essential for maintaining flagellar pocket architecture, with null mutants unable to develop late-stage infections in the sand fly and having reduced virulence in mice [20]. Further, the deletion of FAZ2, a component of the cell body FAZ domain, disrupted the anterior cell tip morphogenesis resulting in FAZ-mediated flagellum-to-flagellum connections [19]. FAZ7B, a FAZ7 paralog which localises to the cell body domain of the FAZ, is also important for cell division and morphogenesis [23]. We have also shown that FAZ34, which localises to the FAZ flagellum domain, is required for flagellum attachment to the *Leishmania* cell body, with FAZ34 deletion resulting in flagellum loss and altered anterior cell tip morphology [22]. Collectively, these findings demonstrate the importance of the FAZ for the morphogenesis of the flagellar pocket and anterior cell tip, which impacts on the pathogenicity and life cycle progression of the parasite.

We have shown that promastigote differentiation to *in vitro* haptomonad-like cells proceeds through five steps from initial adhesion at any point along the flagellum, followed by disassembly of the flagellum and final haptomonad maturation [18]. Furthermore, earlier work and our recent three-dimensional ultrastructural analysis have shown that haptomonad adhesion is mediated by a highly modified, shortened flagellum [18,24].

Ultimately, at the interface of the flagellum and the underlying stomodeal valve, a complex adhesion plaque forms that connects to the surface of the stomodeal valve, with a filamentous network extending from the plaque towards the *Leishmania* cell body and the FAZ. We have identified three kinetoplastid-insect adhesion proteins (KIAPs1-3), which localise to the adhered haptomonad flagellum and are essential for parasite adhesion [15]. Yet the relationship between the cell body FAZ domain and the distal plaque region defined by KIAP1 and KIAP3 is unclear.

Here, we discovered a new set of extra-axonemal flagellar filaments as the flagellum emerged from the cell body and show that KIAP2 is an essential component of them. Deletion of either FAZ2, FAZ5 or FAZ34 impaired *Leishmania* adhesion *in vitro*, and in the latter mutant there was a disconnection of the KIAP2-containing filaments from the anterior cell tip, causing a cell body mispositioning relative to the adhesion plaque and a smaller plaque size with lower levels of KIAPs1 and 3 when compared to the parental cells. Overall, this demonstrates that stable parasite adhesion requires a connection from the cell body through to the adhesion plaque.

## Results

### The FAZ is modified during differentiation to *in vitro* haptomonad-like cells

We have shown by serial electron tomography that there are ultrastructural changes to the anterior cell tip during haptomonad differentiation, including changes to the organisation of the FAZ [18]. To understand the changes in FAZ protein localisation occurring in the FAZ during differentiation to *in vitro* haptomonad-like cells, we generated cell lines expressing mNeonGreen (mNG) and mCherry (mCh)-tagged FAZ proteins characteristic of the different FAZ domains: cell body (FAZ2), intermembrane (FAZ5 and FLA1BP), flagellum (FAZ34), and flagellum exit point (FAZ10), and examined their localisation in cells at different stages of haptomonad differentiation. In the non-adhered promastigotes, all the tagged proteins, except FAZ10, localised as a short line within the flagellar pocket neck region, with FAZ2 and FAZ5 in the cell body and FLA1BP and FAZ34 within the flagellum (Fig 1A, B). FAZ10 localised as a ring at the flagellum exit point from the cell body (Fig S1A). During differentiation to haptomonad-like cells, there was a change to the shape of the fluorescence signal of FAZ2, FAZ5, FLA1BP, and FAZ34 (Fig 1). As differentiation progressed there was an elaboration of the fluorescence signal with additional lobes of signal seen at the anterior cell tip giving the signal from the tagged proteins an L-or Y-shaped appearance (stages iii & iv). During the final stages of differentiation, the line parallel to the flagellar pocket neck shortened until the only observable tagged FAZ protein signal was from the anterior cell tip to give a C-shaped signal in mature (stage v) haptomonad-like cells. In contrast, FAZ10 maintained its ring shape throughout the differentiation process (Fig S1A). These data provide further evidence that the FAZ undergoes structural modifications during haptomonad differentiation.

**Fig. 1.**
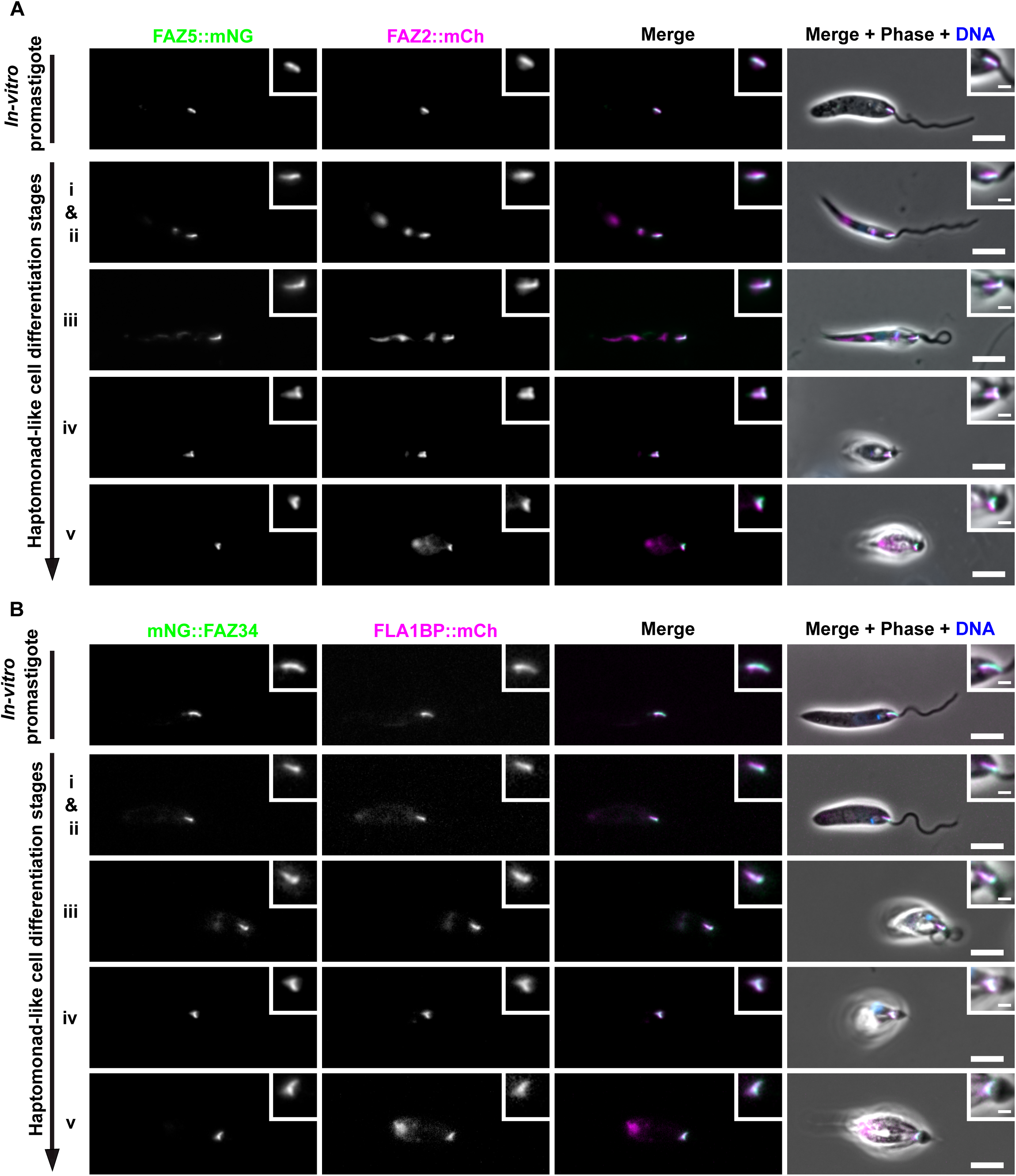
FAZ undergoes structural modification during differentiation of *in vitro L. mexicana* promastigote to haptomonad-like cells. Fluorescence micrographs of *in vitro L. mexicana* promastigote and haptomonad-like cells at different differentiation stages (i-v) expressing mNeonGreen (mNG; green) or mCherry (mCh; magenta) tagged FAZ proteins from the cell body (FAZ2), inter-membrane (FAZ5 and FLA1BP), and flagellum (FAZ34) domains, *in situ*. DNA was stained with Hoechst 33342 (blue). (A) FAZ5 and FAZ2, and (B) FAZ34 and FLA1BP. The insets at the top right corner of each image illustrate the details of the FAZ localisation. The fluorescence intensities were adjusted to show the shape of the FAZ signal. Scale bars: 5 μm; insets: 1 μm.

Next, we examined the positioning of the FAZ relative to KIAPs1-3 using cell lines expressing mNG::FAZ34 and KIAP1::mCh, KIAP2::mCh or mCh::KIAP3. We focussed on FAZ34 because it is found in the flagellum domain of the FAZ [22] and was likely to be positioned closer to the adhesion plaque. In the non-adhered *in vitro* promastigote form, as previously shown [15], KIAP1::mCh localised to the lysosome, with a weak flagellar membrane signal, KIAP2::mCh was positioned within a small region of the flagellum as it emerged from the cell body, while mCh::KIAP3 had a weak cytoplasmic and flagellum signal (Fig S1B). In the *in vitro* haptomonad-like cells, mNG::FAZ34 localised with a C-shaped signal and was positioned adjacent to the enlarged region of the adhered flagellum, with the signals from KIAPs1-3 concentrated in this region (Fig S1C). Notably, FAZ34 localised more closely to KIAP2 than KIAP1 and KIAP3 in the adhered haptomonad-like cells (Fig S1C).

### FAZ2, FAZ5 and FAZ34 are required for haptomonad-like cell adhesion *in vitro*

Given the localisation changes of key FAZ proteins during *in vitro* haptomonad-like cell differentiation, we asked whether these FAZ proteins could be important for haptomonad-like cell adhesion. To investigate this, previously generated FAZ2, FAZ5 and FAZ34 deletion mutants and appropriate add-back cell lines [19,20,22] were allowed to adhere on gridded glass coverslips for 24 hours and the numbers of adhered cells for each cell line compared to those of the parental cells (Fig. 2A, B). The growth rate of the FAZ2, FAZ5 and FAZ34 null mutants as *in vitro* promastigotes was unaffected (Fig. S2A). However, there was a significant reduction in the number of adhered cells for each of the FAZ null mutants, while the add-back of FAZ2 and FAZ34 restored the ability of the parasites to adhere (Fig. 2B). We saw little recovery of adhesion in our initial assays with the FAZ5 add-back cell line (Fig. S2B). In this cell line, the FAZ5 protein was fused to mCh, and we therefore decided to generate a FAZ5 add-back cell line in which the FAZ5 protein was unmodified. We confirmed the presence of the FAZ5 open reading frame by PCR (Fig. S2C). There was an increase in parasite adhesion when using this FAZ5 add-back cell line when compared to the FAZ5 null mutant (Fig. 2B). Together this shows that FAZ2, FAZ5, and FAZ34 are required for adhesion of haptomonad-like cells to substrates.

**Fig. 2.**
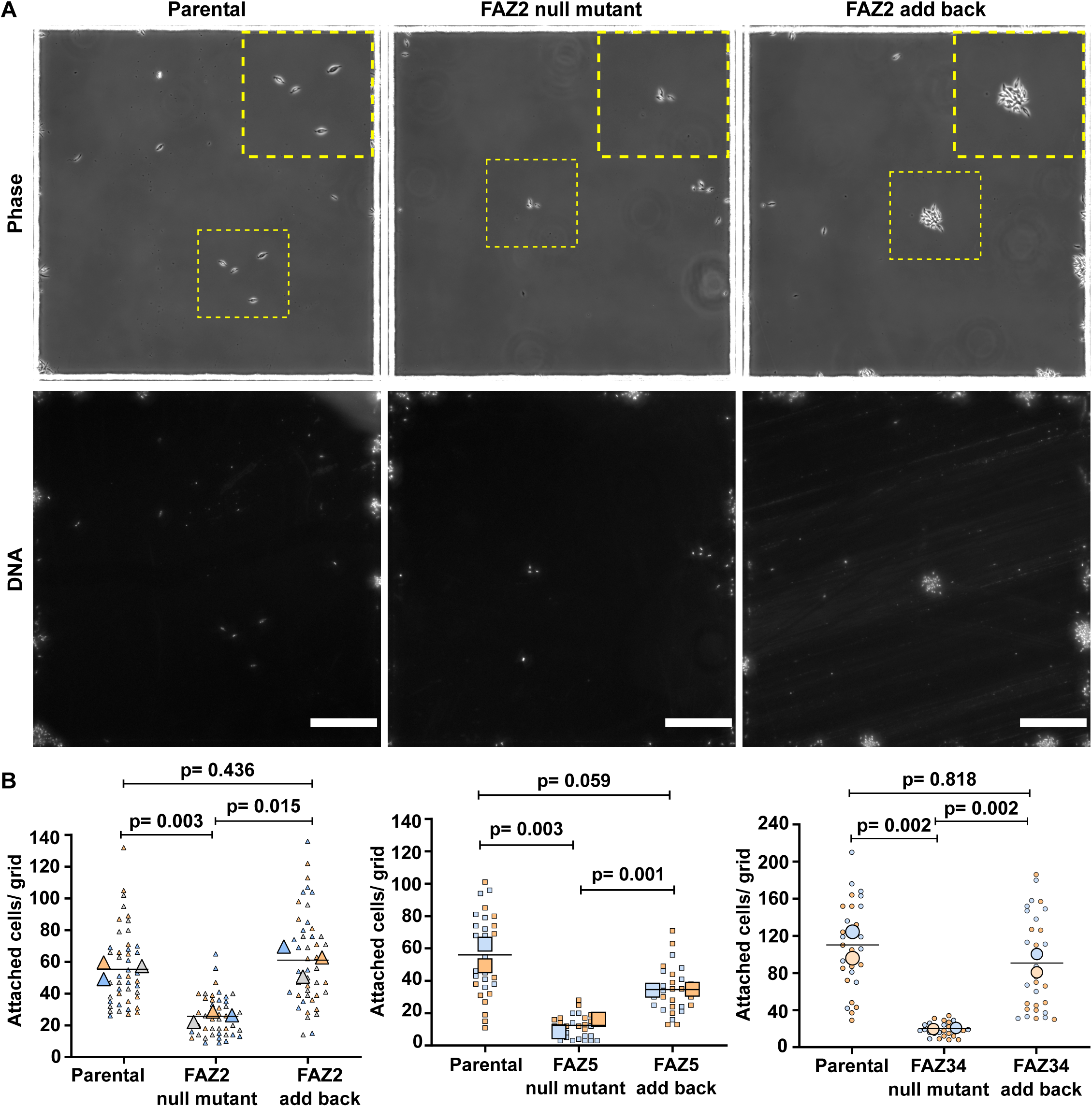
FAZ2, FAZ5 and FAZ34 are required for adhesion of haptomonad-like cells *in vitro*. (A) Phase and Hoechst fluorescence micrographs of *L. mexicana* parental, FAZ2 null mutant and FAZ2 add-back, after 24 h of induced haptomonad differentiation on gridded glass coverslips. Representative images of adhered cells per grid area are shown for FAZ2 parental, null mutant and add-back cell lines. Top: Phase contrast images; bottom: Hoechst 33342-stained DNA. Insets show a magnified view of the yellow-dotted squares, illustrating examples of adhered cells per cell line. Scale bars: 100 μm. (B) Quantification of the number of adhered cells per grid for FAZ2 (left), FAZ5 (middle) and FAZ34 (right) parental, deletion mutants and add-back *L. mexicana* cell lines. The blue, orange and grey colours represent measurements from three independent experiments, for FAZ2, or two independent experiments for FAZ5 and FAZ34. The small triangles, squares or circles indicate counts per grid area while the large ones show the mean values per experiment. The horizontal bars represent the overall mean values of adhered cells per grid area across the experiments. The p-values for FAZ2 and FAZ5 data were calculated using two-tailed Welch’s t-test, while those of FAZ34 data were determined using the Mann-Whitney test.

### FAZ34 is required for the integration of KIAP1 and KIAP3 into the adhesion plaque

Given the position of FAZ34 within the flagellum FAZ domain and its effect on adhesion, we asked whether the loss of FAZ34 impaired the localisation of KIAP1-3 to the adhered flagellum. To address this, we endogenously tagged KIAPs1-3 with mNG in the parental and FAZ34 deletion mutants, allowed the cells to adhere to glass coverslips for 24 h and compared the fluorescence intensities of tagged KIAP1-3 signal in the adhered flagellum of mature parental and FAZ34 null mutant haptomonad-like cells. After adhesion KIAP1, KIAP2, and KIAP3 localised in the expanded region of the flagellum of both the parental and FAZ34 null mutant (Fig. 3A). There was a significantly reduced signal in the FAZ34 null mutant than in the parental cells for both KIAP1 and KIAP3, while KIAP2 signals were similar between the cell lines (Fig. 3A, B).

**Fig. 3.**
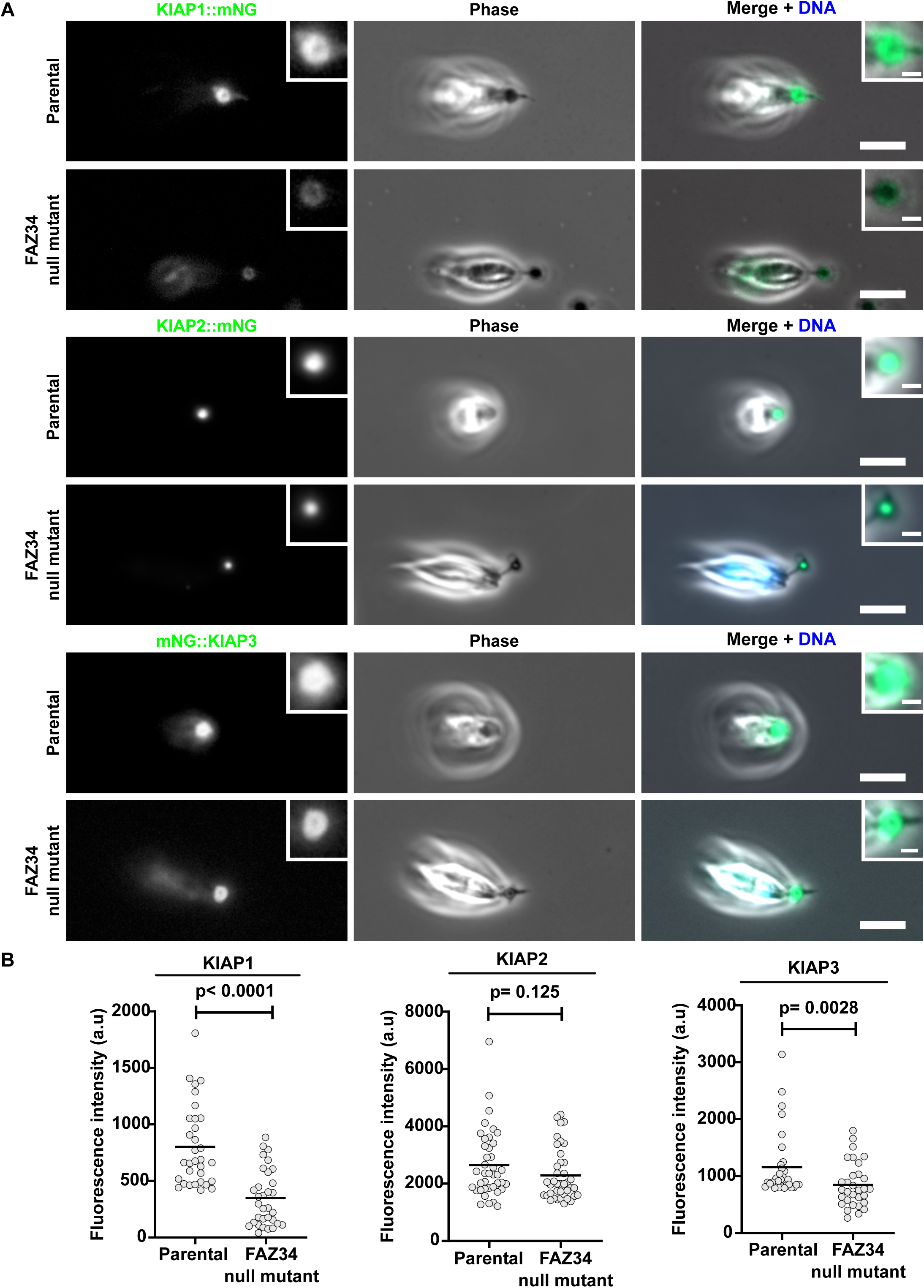
FAZ34 is required for the integration of KIAP1 and KIAP3 into the adhesion plaque. (A) Localisation of mNG-tagged KIAP1, KIAP2, and KIAP3 in parental and FAZ34 null mutant *L. mexicana* haptomonad-like cells. Insets illustrate a magnified view of KIAP1, KIAP2, and KIAP3 in the adhered flagellum. DNA was stained with Hoechst 33342 (blue). Scale bars: 5 μm; inset: 1μm. (C) Quantification of fluorescence intensities of KIAP1 (n = 32 per cell line), KIAP2 (n = 40 per cell line), and KIAP3 (parental: n = 32; FAZ34 null mutant: n = 31) in the adhered flagellum of the parental and FAZ34 null mutant cells after 72 h of adhesion. Fluorescence quantifications were based on average intensity projected z-stack acquired images of 2.25 μm total thickness. The plots show the individual measurements after background correction, with the horizontal bars representing the mean. P-values were calculated using the Mann-Whitney test.

Using super-resolution confocal microscopy, we investigated the effect of the decreased amount of KIAP1 and KIAP3 on the size of the adhesion plaque. We generated parental and FAZ34 null mutant cell lines expressing SMP1 endogenously tagged with mCherry and measured the area of the space bound by the SMP1 signal in mature parental and FAZ34 null mutant haptomonad-like cells. SMP1 is a flagellum membrane marker and is excluded from the adhesion plaque membrane [15]. The area enclosed by the SMP1 signal was significantly smaller in the FAZ34 null mutants compared to the parental cells (Fig S3A, B). Overall, this shows that FAZ34 is required for correct adhesion plaque assembly.

### FAZ34 deletion disconnected KIAP2 and a novel set of filaments from the promastigote anterior cell tip

When we examined our images of mNG-tagged KIAP2 in non-adhered FAZ34 null mutant promastigote cells, we observed a gap separating KIAP2 signal from the anterior cell tip (Fig. 4A). To investigate the changes at the anterior cell tip in the FAZ34 null mutants, we measured the distance between the kinetoplast and the anterior cell tip, as a measure of correct anterior cell tip morphogenesis [19,20,22], and the distance between the kinetoplast and the proximal end of the KIAP2::mNG signal. The distance between the kinetoplast and the anterior cell tip was significantly shorter in the FAZ34 null mutant compared to the parental cells (Fig 4B), as previously observed [22], while the position of the KIAP2::mNG signal did not change relative to the kinetoplast (Fig 4C).

**Fig. 4.**
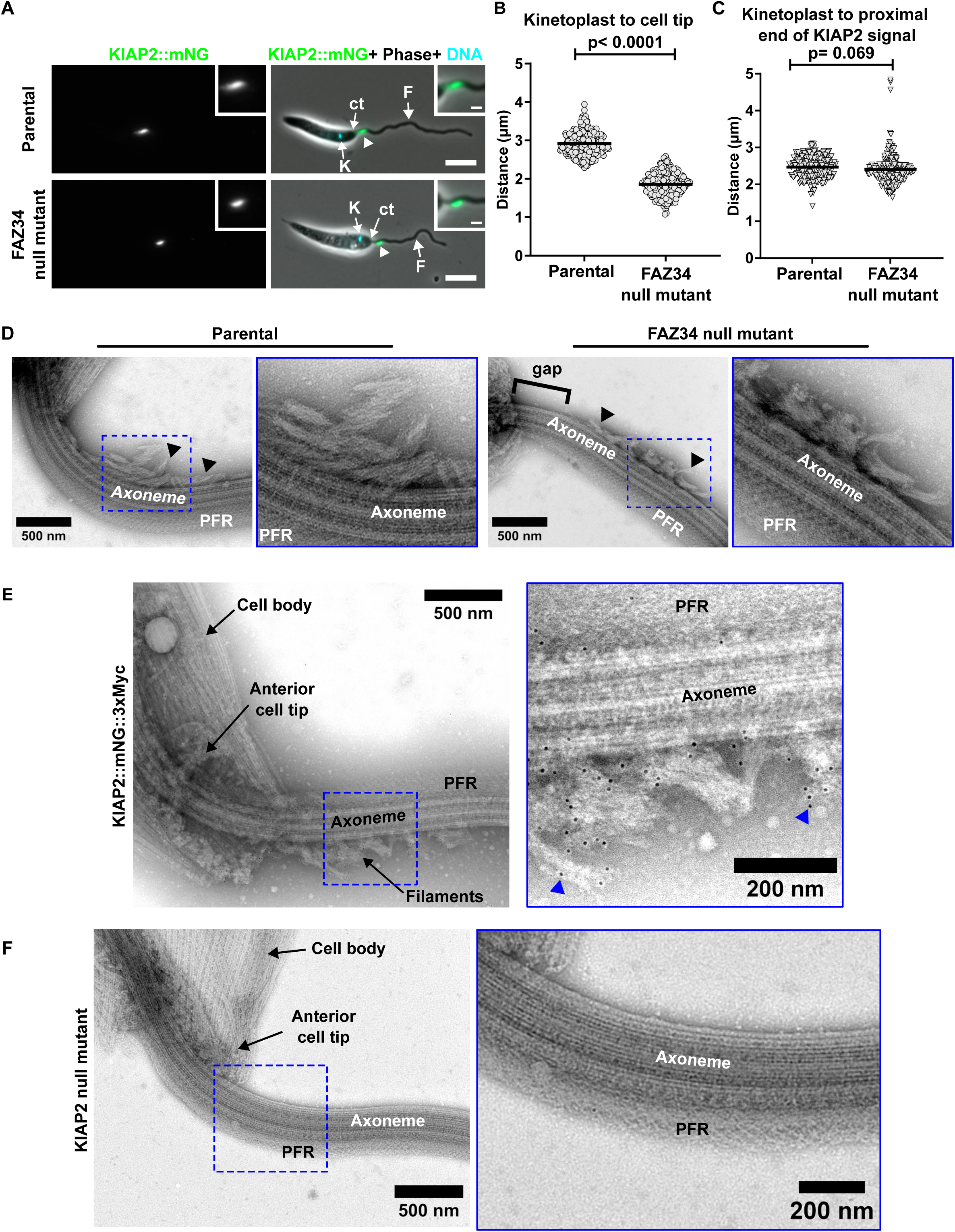
FAZ34 deletion disconnected KIAP2 and a novel set of filaments from the promastigote anterior cell tip. (A) Fluorescence micrographs showing the position of KIAP2 signal (arrowheads) in the parental and FAZ34 null mutant cells. K: Kinetoplast; F: flagellum; ct: cell body tip. Scale bars: 5 μm; inset: 1μm. (B) Quantification of the distance between the centre of the kinetoplast to the anterior cell tip, as a measure of the correct anterior cell tip morphogenesis, in the parental and FAZ34 null mutant cells. Three hundred (n = 300) 1K1N cells were measured for each cell line, and the individual measurements and the mean value (horizontal bar) are shown. (C) Measurement of the position of KIAP2, defined by the distance between the centre of the kinetoplast and the proximal end of KIAP2 signal for the parental and FAZ34 null mutant cells. Data from 200 1K1N cells are plotted, with bar representing the mean. The p-values in B and C were calculated using Welch’s t-test. (D) Negatively stained *L. mexicana* parental and FAZ34 null mutant promastigote cell lines showing the ultrastructure of the filaments (arrowheads) as the flagellum emerges from the cell body. (E) Ultrastructural positioning of KIAP2, by immunogold labelling with 10 nm gold particles. The inset shows a magnified view of the different structures and a high concentration of gold particles (arrowheads) in the filaments. PFR: paraflagellar rod. (F) Negatively stained KIAP2 null mutant whole cell cytoskeleton. KIAP2 deletion resulted in the loss of the filaments.

We asked whether there was a structure responsible for maintaining KIAP2 protein at this specific point in the proximal region of the flagellum. To address this, we examined negatively stained parental and FAZ34 null mutant detergent-extracted cytoskeletons by transmission electron microscopy, enabling us to observe the cytoskeletal structures of the flagellum without the flagellum membrane and cytoplasm. This revealed sets of hitherto undescribed filament bundles on the axoneme as it emerged from the cell body (Fig 4D). The filaments generally lay parallel to the axoneme, on the opposite side to the paraflagellar rod (PFR), a complex, extra-axonemal structure that runs alongside the axoneme [25,26]. In the parental cells the filaments were positioned adjacent to the anterior cell tip, with no clear gap between the anterior cell tip and the filaments, whereas in the FAZ34 null mutant there was a gap (0.42 ± 0.21 µm; n = 46) separating the filaments from the cell tip, which matched our light microscopy observations (Fig 4A).

Given the striking similarity in KIAP2 localisation and the position of these filaments on the flagellum, we investigated KIAP2 localisation at the ultrastructural level by immunogold and electron microscopy, using a cell line expressing KIAP2 endogenously tagged with mNG and 3 Myc epitopes. As a control, we generated a cell line expressing the paraflagellar rod protein 2 (PFR2) fused to the same tag. PFR2 is one of the two most abundant protein components of the PFR [25–27]. Expression of 3Myc-mNG tagged KIAP2 and PFR2 was confirmed by immunofluorescence microscopy (Fig S4A, B). In our electron micrographs of cytoskeletons from cells expressing KIAP2::3Myc::mNG, most gold particles were associated with the filaments, with few or no particles on the PFR and axoneme (Fig 4E). While in the cells expressing PFR2::3Myc::mNG, most gold particles were associated with the PFR, with few or none on the filaments (Fig S4C, D). This shows that KIAP2 is a component of these filaments. Next, we examined detergent-extracted KIAP2 null mutant cytoskeletons by TEM (Fig. 4F). In all cytoskeletons examined (n = 18), we did not observe the filament bundles on the flagellum as it emerged from the cell body. This suggests that KIAP2 is an essential component of these filaments on the flagellum.

In negatively stained dividing cells with a short new flagellum, we observed that the filaments were only present on the old flagellum but not on the new flagellum, suggesting that the assembly of the filaments occurs after the assembly of the microtubule axoneme (Fig S5A). To determine the stage of the *Leishmania* cell cycle at which the filaments appeared, we analysed KIAP2::mNG-tagged promastigote cells by fluorescence light microscopy and classified the cells into different cell cycle stages based on the kinetoplast (K), nucleus (N) and flagellum (F) number [28]. In 1F1K1N cells, we saw a range of KIAP2 signals with a brighter signal in cells with a longer flagellum than those with a shorter flagellum (Fig S5B, C). The KIAP2 signal was weaker in the new flagellum of 1K1N2F cells, with the signal intensity higher in the new flagellum of 2K2N2F cells (Fig S5B). Generally, KIAP2 signal was brighter in the older longer flagella, than in the newer shorter flagella. This confirms that the filaments are assembled into the flagellum after microtubule assembly, with the KIAP2 amount increasing, as the new flagellum elongates.

### Deletion of FAZ34 caused cell body mispositioning relative to the adhesion plaque and KIAP2 in adhered haptomonad-like cells

Given the gap seen between both KIAP2 and the filaments and the promastigote anterior cell tip in the FAZ34 null mutant, we revisited our images of adhered cells of the parental and FAZ34 null mutant to determine whether there was a change in haptomonad-like cell morphology. In the *in vitro* promastigote and early stages of haptomonad-like cell differentiation (i & ii), the flagellum appeared thinner as it emerged from the cell body in all the FAZ34 null mutants examined (n = 100 cells) (Fig. 5A). This thinner region of the flagellum corresponded to the gap separating the KIAP2 signal and the filaments from the anterior cell tip and was likely caused by the disruption of the anterior cell tip morphology (Fig 4A-D; Fig S6). In stably adhered FAZ34 null mutants (stages iii-v), there was a gap separating the cell body from the KIAP2 signal and the adhesion plaque (Fig. 5A; Fig. S6). To characterise the observed phenotype, we acquired z-stack images of mature parental and FAZ34 null mutant haptomonad-like cells, after 24 h of adhesion, and compared the distance between the anterior cell body tip and the centre of the adhesion plaque. This distance was significantly larger (1.51 ± 0.44 µm; n = 25) in the FAZ34 null mutant compared to the parental cells (0.43 ± 0.07 µm; n = 25) (Fig. 5B). However, we did not know whether the increased distance was due to a slower differentiation rate in FAZ34 null mutants versus parental cells. To address this, we allowed the cells to adhere to glass coverslips for 72 h and compared the distances in mature haptomonad-like cells of FAZ34 null mutant and parental cells. The distance between the anterior cell tip and adhesion plaque remained significantly larger (1.37 ± 0.62 µm; n = 70) in the FAZ34 null mutant compared to the parental cells (0.45 ± 0.07 µm; n = 70) and it did not change in comparison to the haptomonad-like cells differentiated for 24 h (Fig. 5B). Overall, all these findings suggest that correct anterior cell tip morphology, mediated by the presence of FAZ34, is required for the positioning of the cell body adjacent to the adhesion plaque during haptomonad differentiation.

**Fig. 5.**
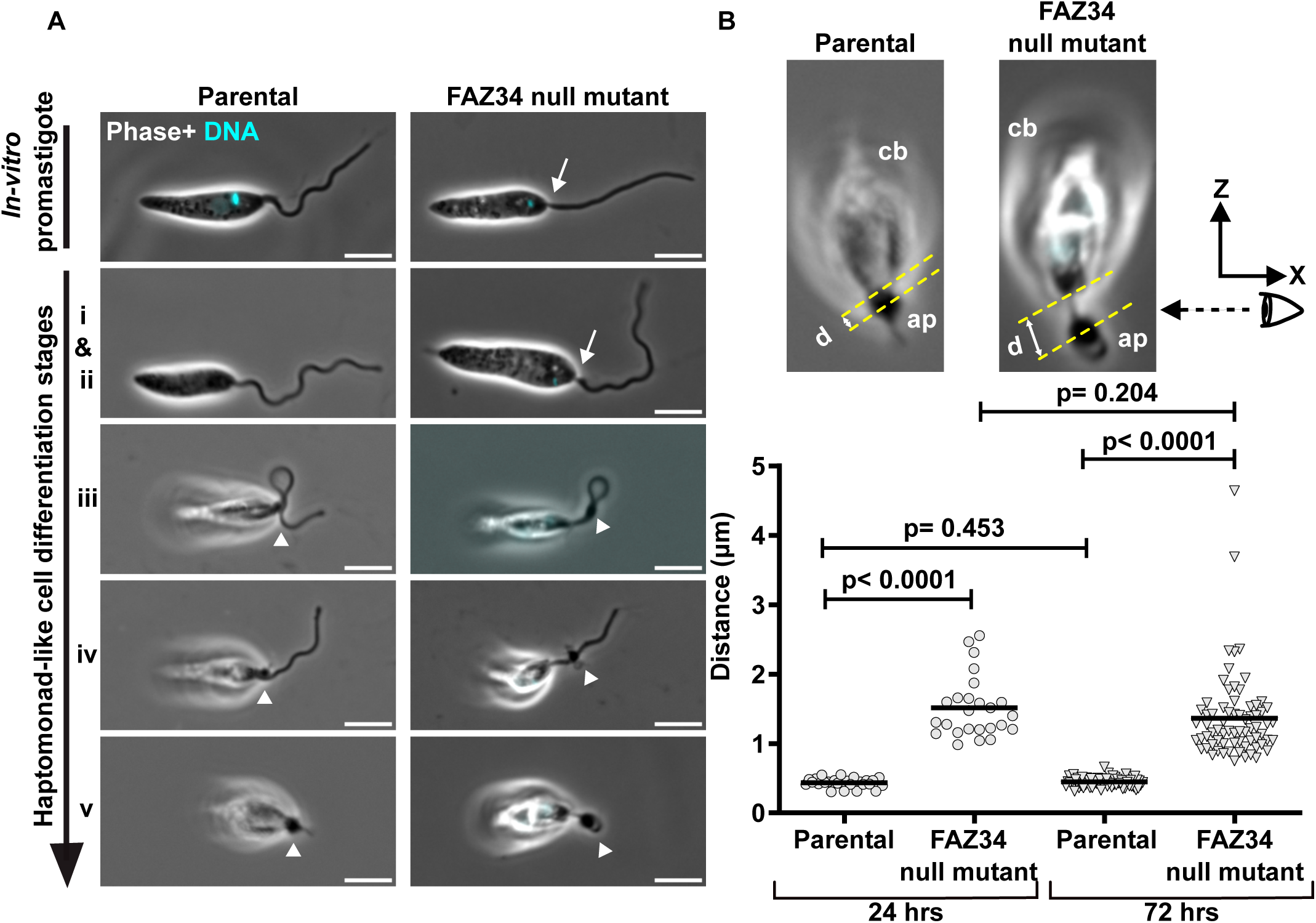
FAZ34 deletion caused cell body mispositioning relative to the adhesion plaque in adhered cells. (A) Phase and Hoechst fluorescence images showing the position of the cell body relative to the adhesion plaque (arrowheads) for the parental and FAZ34 null mutant cells at different haptomonad-like cell differentiation stages (i-v). The arrows show a thinner region of the flagellum as it emerged from the cell body in the FAZ34 null mutant, which was not seen in the parental cells. Scale bars: 5 μm. (B) Measurement of the distance between the anterior cell body tip and the centre of the adhesion plaque for parental and FAZ34 null mutant after 24 h and 72 h of haptomonad-like cell differentiation on glass coverslips. For each cell line, 25 and 70 cells were measured after 24 h and 72 h of differentiation, respectively. The individual measurements (circles or triangles) and the mean (bar) are shown. The p-values were caclulated using two-tailed Welch’s t-test.

## Discussion

*Leishmania* adhesion to the stomodeal valve of the sand fly occurs via a modified flagellum, which contains an adhesion plaque that interacts with the underlying substrate, plus a filamentous network that extends from the plaque and likely connects to the cell body via the FAZ [18]. Strong adhesion requires the progressive disassembly of the long motile flagellum of the promastigote form into a shorter, highly specialised adhesive flagellum of the haptomonad, with accompanying structural and molecular changes to the anterior cell tip [18]. Here, we show that the FAZ undergoes a molecular rearrangement during haptomonad differentiation, with a reorganisation of FAZ proteins from a short linear structure within the flagellar pocket neck region to a C-shaped structure that faces the adhesion plaque. This rearrangement aligns with our previous serial electron tomography data, which showed a reorganisation of the flagellar pocket neck and FAZ region of the cell [18]. The FAZ reorganisation at the anterior cell tip coincided with the enlargement of the flagellum tip, which is associated with the final stages of adhesion plaque assembly and maturation and is likely required to form a strong cytoskeletal connection extending from the cell body to the adhesion plaque.

The FAZ is important for anterior cell tip and flagellar pocket morphogenesis in *Leishmania*, with the loss of FAZ proteins associated with a reduction in the kinetoplast to cell tip distance [19,20,22]. Moreover, FAZ5 deletion resulted in the loss of the distinctive asymmetric cell tip, where the cell body extends further along the side of the flagellum associated with the FAZ [20], while the loss of FAZ2 resulted in an extension of this asymmetry with the cell body extending further along the flagellum [19]. The FAZ is a complex cytoskeletal structure, with distinct cell body, intermembrane, and flagellum domains. Deletion of proteins from each of these domains (FAZ2 – cell body; FAZ5 – intermembrane; FAZ34 – flagellum) severely impaired the adhesion ability of *Leishmania* parasites *in vitro* that was restored in the add-back cell lines. Moreover, in the FAZ34 null mutant, there were defects in adhesion plaque assembly, with reduced levels of KIAP1 and KIAP3 in the adhesion plaque and a reduced plaque size. These defects likely translate to a weaker connection to the underlying substrate, reflected in the reduced number of stably adhered parasites (model in Figure 6). Overall, this demonstrates the importance of the FAZ for the establishment of strong and stable adhesion to a substrate, with disruption of anterior cell tip morphogenesis likely causing defects in adhesion plaque assembly.

**Fig. 6.**
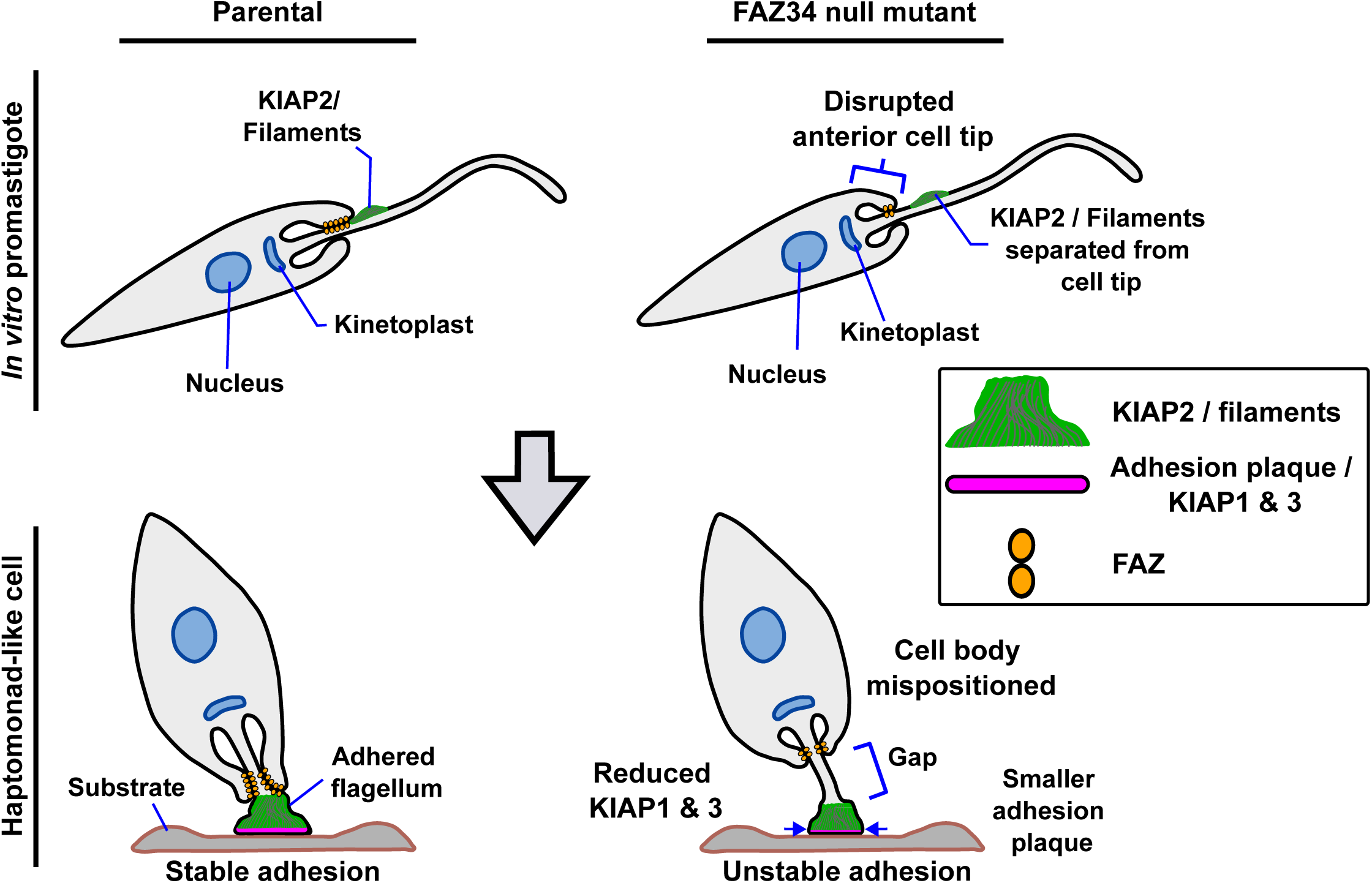
Model of the roles of FAZ in *Leishmania* adhesion. A schematic of the parental and FAZ34 null mutant promastigote and haptomonad-like cells based on *in vitro* observations. FAZ34 deletion disrupts the anterior cell tip morphology, resulting in the separation of KIAP2 and the flagellar filaments from the anterior cell tip in the promastigote and haptomonad-like cells, with the cell tip positioned further from the adhesion plaque in the latter when compared to parental cells. This is associated with defects in adhesion plaque assembly, with lower levels of KIAP1 and KIAP3 in the plaque and a reduced plaque size, which translates to unstable adhesion in the FAZ34 null mutant.

The localisation of KIAP2 to a region of the flagellum in promastigotes as it emerges from the cell body prompted us to investigate the ultrastructure of this region by TEM of whole mount negatively stained cytoskeletons. We identified a novel set of filament bundles in this much studied stage of the life cycle that lay parallel to the axoneme, on the opposite side to the PFR and adjacent to the anterior cell tip. KIAP2 localisation to these filaments was confirmed by immunogold staining and its presence was required for their assembly. KIAP2 encodes a calpain-like protease domain that lacks the catalytic residues, and this domain is thought to mediate protein-protein interactions. It is likely that KIAP2 is the main component of these novel filaments; however, its filament-forming capabilities have yet to be investigated. Alternatively, KIAP2 may be important for stabilising filaments formed by an additional as yet unidentified protein. We have shown that in the modified flagellum of haptomonads there is a filamentous network extending from the adhesion plaque to the cell body, with KIAP2 potentially localised along these filaments [15,18]. The presence of KIAP2-containing filaments in the proximal region of the promastigote flagellum and the modified haptomonad flagellum indicates that these filaments will likely have common constituents. Given that KIAP2 is required for both parasite adhesion [15] and filament formation, this suggests these filaments are also central to *Leishmania* adhesion.

In dividing cells, we saw that the KIAP2 signal was initially absent in the new flagellum, with longer flagella having a brighter signal. This shows that these filaments are assembled into the flagellum after microtubule axoneme assembly, similar to the PFR [26,29]. Moreover, the post-axonemal assembly of KIAP2 will likely impact the adhesive capacity of the daughter cell that inherits the new flagellum. One might envisage that a cell inheriting the new flagellum, with less KIAP2, will be less likely to adhere at the start of the cell cycle than later when the amount of KIAP2 in the flagellum has increased.

In the FAZ34 null mutant promastigotes, the close association between the KIAP2, flagellar filaments and the anterior cell tip was lost, resulting in a separation between the KIAP2, flagellar filaments and anterior cell tip (Figure 6). This separation was also seen in the adhered FAZ34 null mutants, with the cell tip positioned further from the adhesion plaque when compared to parental cells. The separation between the cell tip and the adhesion plaque observed in adhered FAZ34 null mutant is likely due to the disruption of the anterior cell tip morphology caused by the loss of FAZ34 [22]. This suggests that the close association of KIAP2 and the flagellar filaments with the anterior cell tip is required for stable parasite adhesion and is mediated by the FAZ.

These flagellar filaments have not been explicitly described in *Leishmania* before; however, when we revisited our collection of thin-section transmission electron micrographs of this region of the cell, we often saw a bulge in the flagellum as it emerged from the cell body. In this bulge, there was little pattern to the electron density, so we suspect that these filaments are not well-preserved using traditional EM fixatives and methodologies. The bulge is also found on the side of the flagellum associated with the FAZ (Fig S7), further supporting the potential connections between these flagellar filaments and the FAZ. Interestingly, filaments have been observed in the flagellum of *Crithidia fasciculata* in a similar position and *Paratrypanosoma confusum* has a large bulge in its flagellum as it emerges from the cell body [31,32]. In the latter case, this large bulge is associated with the adhesion of this parasite to a substrate. This suggests that these flagellar filaments are an ancestral feature of the kinetoplastid parasites that adopt a liberform morphology [33] and are likely important for their adhesion to a range of substrates.

Furthermore, the presence of these filaments in the flagellum of motile stages indicates that these parasites are primed for adhesion, with molecular components already present, enabling the efficient establishment of a stable connection to a substrate.

In conclusion, we have identified and initially characterised a new set of flagellar filaments, which contain the adhesion protein KIAP2, and likely act as a coordinating foundational structure for parasite adhesion. While also showing the importance of maintaining a connection from adhesion plaque through to the cell body for stable *Leishmania* adhesion to a substrate.

## Resource availability Lead contact

Requests for further information, resources and reagents should be directed to and will be fulfilled by the lead contact, Jack Sunter

## Materials availability

This study did not generate new unique reagents.

## Data and code availability

Any additional information required to reanalyse the data reported in this paper is available from the lead contact upon request.

## Supporting information

Supplemental information titles and legends

## Acknowledgements

BOO was funded by the Nigel Groome PhD studentship and the Sunter lab is funded by the Wellcome Trust. We thank Dr Clare Halliday (Oxford Brookes University) and Ed Rea (Oxford Brookes University) for providing electron micrographs and Dr Jorge Arias Del Angel (University of Würzburg) for assistance with developing scripts for area measurements. We thank Dr Richard Wheeler (University of Edinburgh), Dr Uli Dobramysl (University of Oxford), Prof Keith Gull (University of Oxford) for many discussions. We also acknowledge the expertise of the Oxford Brookes University Centre for Bioimaging.

## Author contributions

Conceptualisation: JDS

Formal analysis: BOO

Funding acquisition: BOO, JDS

Investigation: BOO, RY, AOM, FML, SV, JDS

Methodology: BOO, RY, AOM, FML, SV, JDS

Project administration: BOO, JDS

Supervision: RY, SV, JDS

Validation BOO, RY, AOM, FML, SV, JDS

Visualisation: BOO

Writing (original draft): BOO, JDS

Review and editing: BOO, RY, AOM, FML, SV, JDS

## Supplemental information titles and legends

Document S1. Figures S1-S7

## Methods

### Cell culture

*Leishmania mexicana* promastigotes (WHO strain MNYC/BZ/62/M379), expressing Cas9 nuclease and T7 polymerase were grown at 28[C in M199 medium with Earle’s salts and L-glutamine (Life Technologies), 26 mM NaHCO_3_, 5 µg/mL haemin, 40 mM HEPES-HCL (pH 7.4) and 10% foetal bovine serum (FBS). The cells were maintained in the logarithmic growth phase by regular sub culturing.

### Generation of FAZ and KIAPs 1-3 tagging, deletion and add-back constructs

Constructs and single guide RNAs (sgRNAs) for endogenous mNeonGreen or mCherry tagging of *L. mexicana* FAZ, PFR2, SMP1 and KIAPs 1-3 genes were generated by PCR method, according to Dean *et al.,* [34] and Beneke *et al.,* [35], using pPLOT plasmids with either blasticidin, puromycin or neomycin resistance genes as the templates, or pLPOT plasmid with blasticidin resistance gene as the template [30]. The FAZ gene deletion constructs were generated by PCR using two pT plasmids with different resistance genes (blasticidin, puromycin or neomycin) and the G00 as the template [35]. Primers and sgRNAs for the tagging and deletion constructs were designed using LeishGEdit (http://www.leishGEdit.net) [35]. FAZ5 add-back construct was generated as previously described [20,36]. All the transfections were performed using the X-001 programme on the Amaxa Nucleofector II device and successful transfectants selected, after 6 hours, using blasticidin (5 µg/ml), puromycin (20 µg/ml) or G-418 (20 µg/ml) (Melford Laboratories).

### Growth curves

*Leishmania mexicana* promastigotes were set up at 1 x 10^5^ cells/ mL in M199 supplemented with 10% FBS. Cumulative cell growth was measured daily by counting the number of cells using the Beckman Coulter Counter.

### *In vitro* haptomonad differentiation assays

Axenic haptomonad cells were generated according to Yanase *et al*.,[18]. Briefly, gridded glass coverslips grid-500, (ibidi; 10816) were cut into ∼5 x 5 mm pieces and sterilised with 100% ethanol for 5 min in a sterile Petri dish. The coverslips were washed twice with 5 ml of M199 media, transferred to a 24-well plate containing 1 mL of M199 medium, and washed two more times with 1 mL of M199. Following the final wash, promastigotes at 5×10^6^ cells/mL of M199 medium were added to the coverslips and allowed to differentiate for 24 h at 28 [C with 5% CO_2_.

### Quantitative analysis of the effect of FAZ deletion on haptomonad adhesion, *in vitro*

Gridded glass coverslips grid-500 (ibidi; 10816) were prepared as described above and three coverslips were used for each cell line. Log phase promastigotes of FAZ2, FAZ5 and FAZ34 parental, deletion mutants, and add-back cell lines (5×10^6^ cells/mL) were grown on the ∼5 x 5 mm pieces in a 24-well plate with 1 mL of M199 medium for 24 h at 28 [C with 5% CO_2_. The coverslips were washed twice by transferring them to wells with 1 mL DMEM, and then incubated with 1 mL DMEM containing 1 µg/mL of Hoechst 33342 for 5 min. The coverslips were then washed twice with 1 mL DMEM and mounted, with the top side facing up, on a super frost microscope slide with premarked circular wells, created using the ImmEdge hydrophobic barrier pen (Vector laboratories; H-4000). Another glass coverslip was mounted carefully onto the gridded coverslips to avoid air bubbles. Adhered cells were imaged in a 500 µm x 500 µm grid area using the Zeiss Axio ImagerZ2 upright microscope with the Zeiss Plan-Apochromat 20x/0.8NA PH2 objective and Hamamatsu Flash 4 camera (Hamamatsu Photonics, Hamamatsu, Japan). Images of adhered cells in five different grids were acquired for each coverslip using the phase (5 ms exposure, TL-VIS LED Lamp illumination) and H3342 (Zeiss filter set 49; 50 ms exposure time, Zeiss HBO100 mercury bulb illumination using Osram HBO103/W2 bulbs) channels in ZEN 2.3 Blue (release version 69.1) software. Adhered cells in each grid area were manually counted in Fiji [37], using the phase contrast image and Hoechst signal.

### Widefield microscopy

For live cell imaging of tagged or untagged *L. mexicana* promastigotes, 1 mL of log phase cell cultures (∼1×10^7^ cells/mL) were washed twice with 1 mL of DMEM. The cells were resuspended in 1 mL DMEM containing 1 µg/mL of Hoechst 33342 and incubated at RT for 5 min. Following incubation, cells were washed twice with 1 mL of DMEM and resuspended in a 100 µL final volume. One microliter (1 µL) of the cell suspension was added to the centre of premarked circular wells on a super frost microscope slide. The cells were mounted with a glass coverslip and imaged using the Zeiss Axio ImagerZ2 upright microscope with the Zeiss 63x/1.4NA PH3 oil-immersion Plan-Apochromat objective and Hamamatsu Flash 4 camera. For KIAP2 signal intensity observation in dividing cells, we imaged the cells using the Zeiss alpha Plan-Apochromat 100x/1.46NA PH3 oil immersion (UV) M27 objective. The images were acquired in ZEN 2.3 Blue software using the phase (5 ms exposure), mCherry (Zeiss filter set 43 HE; 150 ms exposure, Zeiss HBO100 mercury bulb illumination using Osram HBO103/W2 bulbs), EGFP (Zeiss filter set 38 HE; 150 ms exposure, Zeiss HBO100 mercury bulb illumination using Osram HBO103/W2 bulbs) and H3342 (60 ms) channels. We performed all washes by centrifuging cells at 800xg for 3 min.

For the haptomonads, 18 x 18 mm high-performance coverslips thickness No. 1.5H (Zeiss) were sterilised with 100% ethanol for 5 min in a sterile Petri dish, washed twice with 5 ml of M199 media and transferred to 6-well plates. Further washes were performed twice with 2 mL M199. Promastigotes at 5×10^6^ cells/mL in 2 mL of M199 medium were added to the coverslips and allowed to differentiate at 28 [C with 5% CO_2_ for 24 h (for localisation and intensity measurements) or 72 h with the medium being changed every 24 h (for cell body mispositioning assays). The coverslips were washed twice with 2 mL of DMEM, incubated for 5 min with 2 mL DMEM containing 1 µg/mL of Hoechst 33342 and washed twice with 2 mL of DMEM. Washed coverslips were mounted onto a super frost microscope slide with premarked square wells and another coverslip mounted on top. We imaged the adhered live cells using the Zeiss Axio ImagerZ2 upright microscope with the Zeiss 63x/1.4NA PH3 oil-immersion Plan-Apochromat objective and Hamamatsu Flash 4 camera. Z-stacks were acquired on the phase (5 ms), mCherry (150 ms), EGFP (150 ms) and H3342 (60 ms) channels using a voxel size of 0.103 µm x 0.103 µm x 0.25 µm. All the z-stack images were acquired in ZEN 2.3 Blue software and analysed using Fiji [37]. Quantification of fluorescence intensities of KIAPs in the adhered flagella was based on average intensity projection using nine z-stack slices of 2.25 μm total thickness. Same analysis settings were applied for each comparison group and the figures generated using the QuickFigures plugin in Fiji [37,38]

### Confocal microscopy

For the confocal microscopy of adhered live cells, 1×10^6^ cells/mL log phase promastigotes were grown in 2 mL of M199 in the ibidi’s μ-dish 35 mm (high glass bottom) culture dishes for 24 h at 28 [C with 5% CO_2_. To remove non-adherent cells, the dishes were washed more than five times with fresh M199, prewarmed at RT. The cells were imaged using an inverted Zeiss LSM880 with Airyscan, 100x oil-immersion alpha Plan-Apochromat objective (1.46NA; DIC M27 Elyra), 561nm laser and Physik Instrumente P-737 piezo stage. We acquired confocal z-stack images in super-resolution Airyscan mode using line scanning and the settings previously described [15]. The images were processed in ZEN 2.3 SP1 FP3 Black (release version 14.0) software and analysed using Fiji [37]. For the quantification of the adhesion plaque size, nine Airyscan-processed z-stack slices of 0.45 μm total thickness were subjected to average intensity projection. These were then binarised in MATLAB (R2023a) and a pixel intensity threshold of 0.5 used to determine the boundaries of the area enclosed by the SMP1 signal. To exclude cell debris, images with a circularity lower than 2 were selected.

### Immunofluorescence microscopy

Approximately 1×10^7^ log phase promastigotes of C-terminally tagged KIAP2 and N-terminally tagged PFR2 cell lines expressing mNeonGreen fused to 3myc epitope (KIAP2:: 3Myc::mNG and 3Myc::mNG::PFR2) were washed three times with 1 ml of Voorheis modified PBS (vPBS) [39]. The cells were resuspended in 100 µL of vPBS and 50 µL of the cell suspension was settled in the pre-marked circular wells on a super frost microscope slide (Thermo Scientific). The parasites were allowed to adhere to the slides for 5 min, then fixed for 10 min with 4% paraformaldehyde (PFA) in vPBS at RT. Unreacted PFA was quenched with 1% glycine in PBS for 5 min at RT. Whole-cell cytoskeletons of adhered KIAP2::mNG::3Myc and mNG::3Myc::PFR2 promastigotes were prepared by permeabilising the cells with 1% NP-40 in PEME for 30 sec. The cells were fixed with methanol for 20 min at −20 [C, washed with PBS for 20 min at RT, and blocked with 1% BSA in PBS for 1 h at RT. The cells were then incubated with 1:200 c-Myc monoclonal antibody (9E10; MA1-980) in PBS containing 1% BSA for 1 h at RT. The cells were washed four times, with 5 min incubation during each wash, and probed with Alexa Fluor 546 goat anti-mouse secondary antibody (Invitrogen) diluted at 1:1000 in 1% BSA in PBS for 1 h at RT. The slides were washed twice with PBS for 5 min each, incubated with 1 µg/mL of Hoechst 33342 in PBS for 5 min, washed once with PBS for 5 min, and mounted with Vectashield mounting medium for imaging. The images were acquired under the phase, mCherry, EGFP and H3342 channels at 63x magnification, as described above.

### Negative staining

1×10^7^ log phase promastigotes of parental and FAZ34 null mutant cell lines were settled on glow-discharged carbon and formvar-coated nickel grids (200 mesh; Agar Scientific), mounted on tweezers, for 2 min at RT. Whole cytoskeletons were prepared by inverting the grids onto ∼1 mL drop of 1% IGEPAL CA-630 (Sigma; I3021) in PEME for 5 min at RT. The samples were rinsed twice with ∼1 mL drops of PEME and fixed with 2.5% glutaraldehyde (EM grade) in PEME for 10 min at RT. Fixed samples were washed twice in drops of PEME (5 min/ wash), transferred to ∼1 mL drop of ddH_2_O and stained with 1% aurothioglucose in ddH_2_O. The grids were allowed to dry, followed by imaging on a Jeol JEM-1400 Flash transmission electron microscope operating at 120 kV. The images were acquired using a OneView 16-megapixel camera (Gatan/Ametek version 3.52.3932.0, Pleasanton, CA).

### Immunogold labelling

Whole-cell cytoskeletons of approximately 1×10^7^ log phase promastigotes of the parental cells and cell lines expressing KIAP2::mNG and mNG::PFR fused to 3myc epitope (KIAP2::mNG::3Myc and mNG::3Myc::PFR2) were prepared as described above. The samples were fixed with 4% PFA in PEME for 10 min at RT and the unreacted PFA was quenched with 1% glycine in PEME for 5 min at RT. Each grid was transferred to a drop of PEME (50 µL) for 5 min and blocked for 30 min in a 50 µL drop of 1% BSA in PEME. The samples were incubated with c-Myc monoclonal antibody (9E10; MA1-980) diluted at 1:20 in the blocking buffer for 1 h at RT, followed by three consecutive washes in 50 µL drops of the blocking buffer (5 min/wash).

Washed samples were probed with goat anti-mouse secondary antibody conjugated to 10 nm gold (Sigma; G7652; 1:50 dilution in the blocking buffer) for 1 h at RT. The grids were then washed by transferring them to three consecutive drops of PEME (∼1 mL/drop; 5 min/wash) and fixed with 2.5% glutaraldehyde in PEME for 10 min at RT. Fixed samples were washed twice in drops of PEME (5 min/ wash), and negatively stained with 1% aurothioglucose in ddH_2_O. The samples were allowed to dry and imaged on a Jeol JEM-1400 Flash transmission electron microscope operating at 120 kV, equipped with a OneView 16-megapixel camera (Gatan/Ametek version 3.52.3932.0, Pleasanton, CA).

### Statistical analysis

The means were calculated in Microsoft Excel. Statistical differences between groups were estimated using the two-tailed Welch’s *t*-test or the Mann-Whitney test at p< 0.05 significance level in GraphPad Prism (v9.5). Data plots were created using GraphPad Prism.

