## Supplemental information titles and legends for "Discovery of a novel flagellar filament system underpinning *Leishmania* adhesion to surfaces"

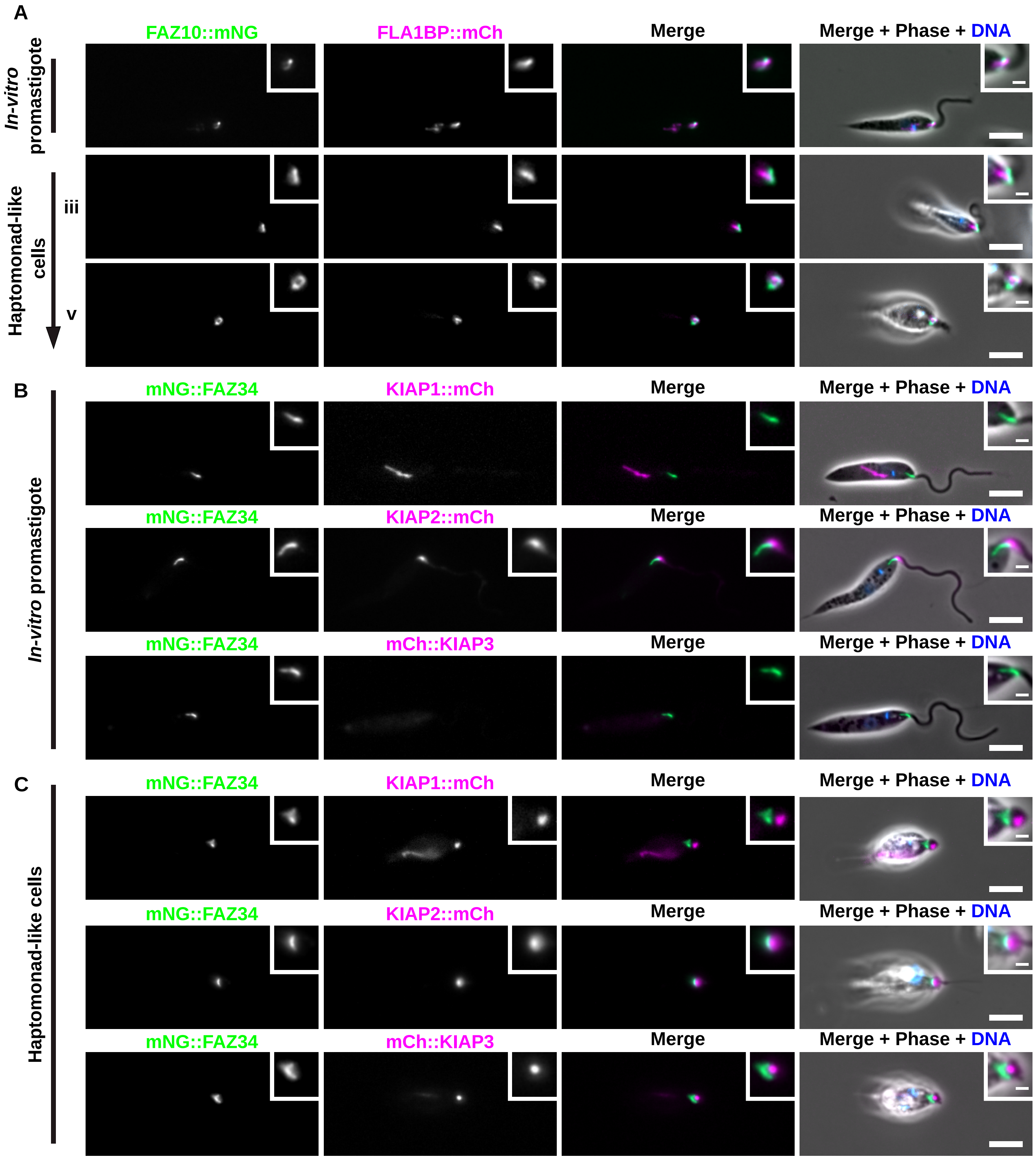


**Fig S1. Localisation of FAZ and KIAPs1-3 in *L. mexicana in vitro* promastigote and haptomonad-like cells.** (A) Fluorescence micrographs of *L. mexicana* promastigote and haptomonad-like cells expressing mNeonGreen (mNG) and mCherry (mCh) tagged FAZ10 and FLA1BP proteins at stages iii and v of differentiation. Insets illustrate a magnified view of the FAZ signal at the different stages. (B-C ) Localisation of mNG-tagged FAZ34 and mCherry-tagged KIAP1, KIAP2 or KIAP3 in *L. mexicana* promastigote (A) and haptomonad-like cells (B). Insets show a magnified view of FAZ34 and KIAPs1-3 signals and the position of FAZ34 relative to KIAPs1-3 in promastigotes and haptomonad-like cells. The nucleus and kinetoplast DNA are shown in blue, following Hoechst 33342 staining. Scale bars: 5 μm, insets: 1 μm.

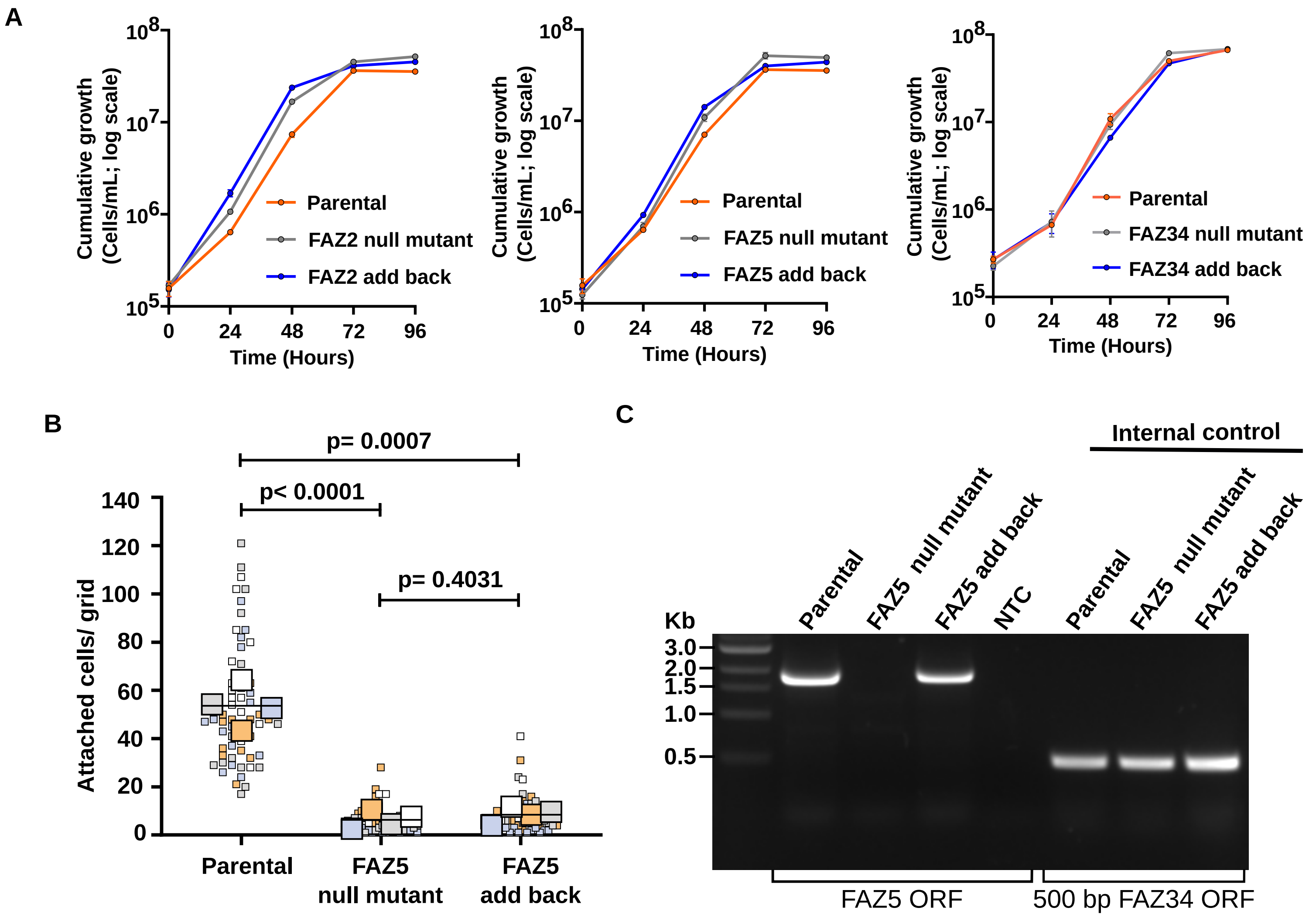


**Fig S2.** (A) Cumulative growth curves of parental and FAZ2 (left), FAZ5 (middle), and FAZ34 (right) null mutant and add-back cell lines. Data from three independent experiments are shown with the error bars representing the mean ± SD. (B) Quantification of the number of adhered haptomonad-like cells for the parental, FAZ5 null mutant and an add-back cell line in which FAZ5 protein was fused to mCherry. The orange, blue, white and grey colour codes represent measurements from four independent experiments (n = 4), with the small squares indicating individual counts per grid area while the large squares show the average counts per experiment. The horizontal bars represent the overall mean value across the four indipendent experiments. The p-values were calculated using the Welch's two-tailed t-test. (C) Confirmation of deletion and add-back of FAZ5 gene. Genomic DNA from the parental, FAZ5 null mutant and FAZ5 add-back cell lines were analysed by PCR and resolved on 1% agarose gel. We observed a band of ~1.9 Kb, corresponding to the FAZ5 ORF, in the parental and FAZ5 add-back, but not in the FAZ5 null mutant cell line, confirming the successful deletion and add-back of the gene. The successful amplification of ~500 bp FAZ34 internal control in all the cell lines indicated a good quality DNA and efficient PCR amplification. NTC: nuclease-free water

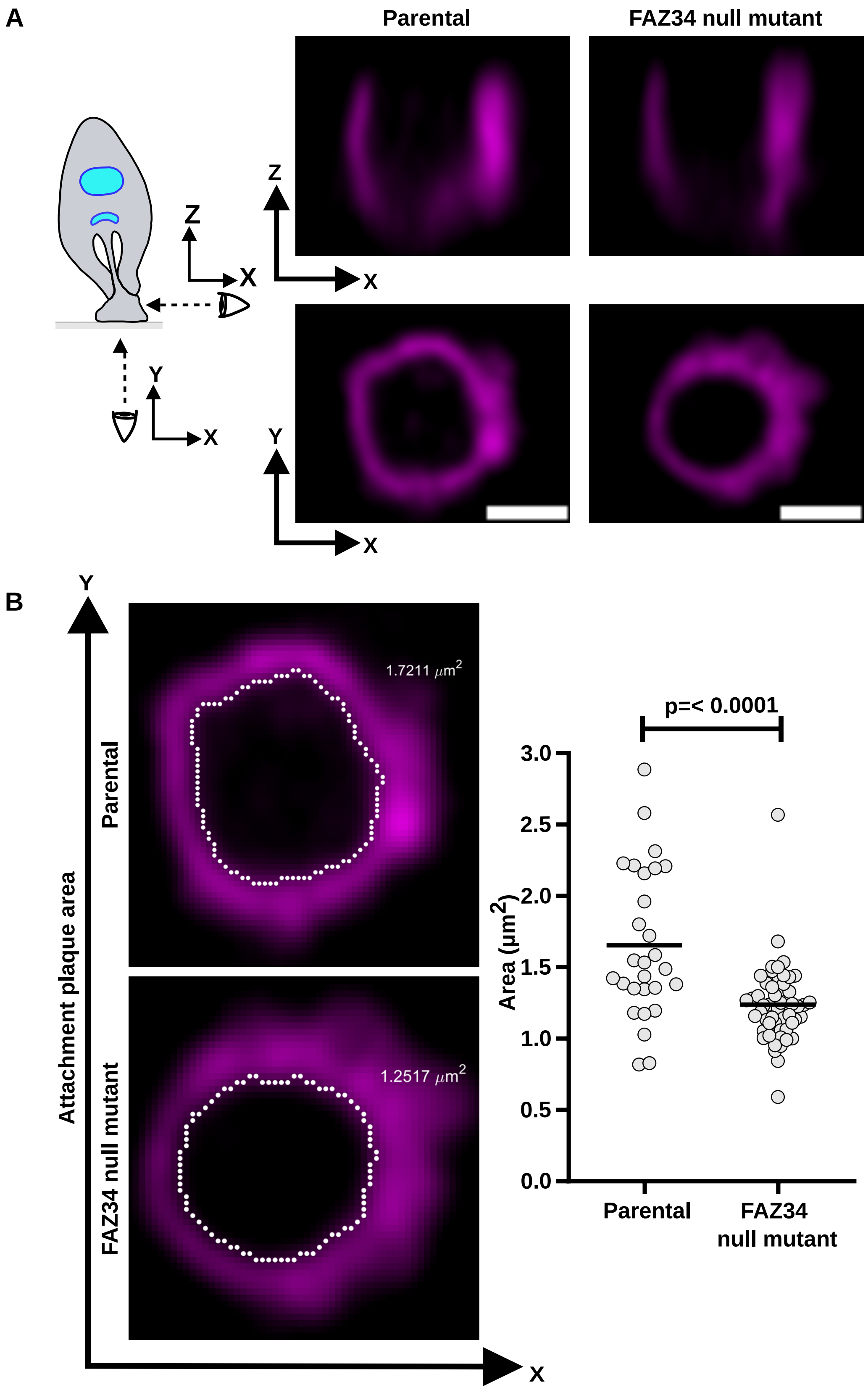


**Fig S3. FAZ34 deletion reduced the size of the adhesion plaque.** (A) Z-stack confocal microscopy analysis of the adhered flagella of parental and FAZ34 null mutant cells expressing SMP1 endogenously tagged with mCherry. The schematic diagram illustrates the different viewing positions: X-Y (bottom view), and X-Z (side view). Scale bars: 1 μm. (B) Quantification of the area enclosed by the SMP1 signal in the parental (n = 28) and FAZ34 null mutant (n = 54) cells after 72 h of adhesion on glass coverslips. For the area measurements, we binarised the Airyscan-processed images of 0.45 μm thickness in MATLAB (R2023a) and used a 0.5-pixel intensity threshold to determine the boundaries of the area enclosed by the SMP1 signal. To exclude cell debris, images with a circularity lower than 2 were selected. We calculated the p-values using the Mann-Whitney test.

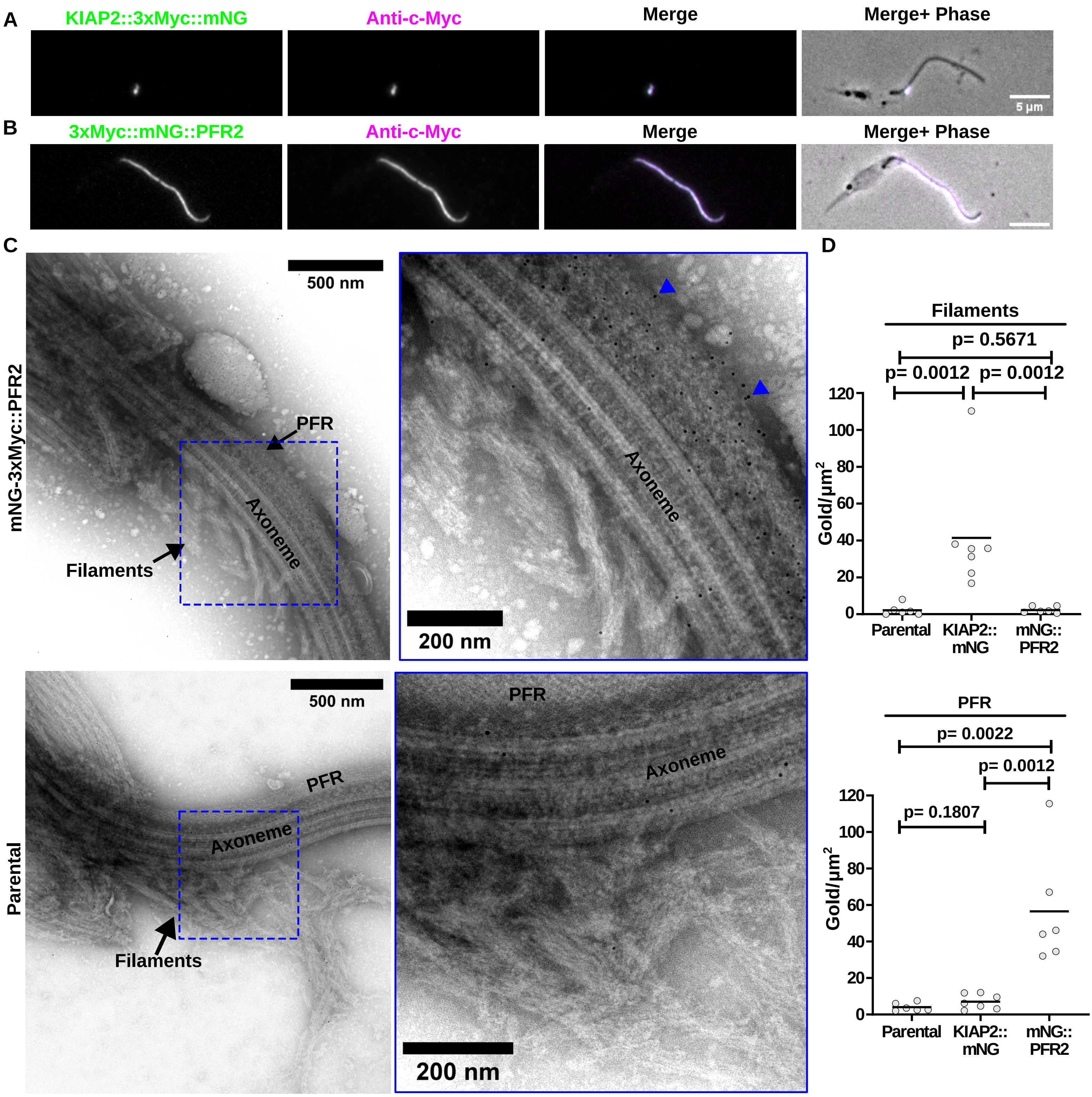

A
KIAP2::3xMyc::mNG
Anti-c-Myc
Merge
Merge+ Phase
5 µm
B
3xMyc::mNG::PFR2
Anti-c-Myc
Merge
Merge+ Phase
C
mNG-3xMyc::PFR2
500 nm
PFR
Axoneme
Filaments
200 nm
Axoneme
Parental
500 nm
PFR
Axoneme
Filaments
PFR
Axoneme
200 nm
D
Filaments
p= 0.5671
p= 0.0012
p= 0.0012
Gold/µm2
Parental
KIAP2:: mNG
mNG:: PFR2
PFR
p= 0.0022
p= 0.0012
p= 0.1807
Gold/µm2
Parental
KIAP2:: mNG
mNG:: PFR2

**Fig S4. FAZ34 deletion separated KIAP2 and a novel set of filaments from the anterior cell tip in *L. mexicana in vitro* promastigote.** (A, B) Confirmation of 3Myc::mNG tagged KIAP2 and PFR2 expression in whole mount cytoskeletons of *L. mexicana* promastigotes by immunofluorescence. (A) Fluorescence micrographs showing the expression of a C-terminally tagged KIAP2 (KIAP2::3Myc::mNG) as the flagellum emerged from the cell body and, (B) N-terminally tagged PFR2 (3Myc::mNG::PFR2). Detergent extracted whole cell cytoskeletons were prepared for immunofluorescence microscopy using c-Myc monoclonal primary antibody (9E10; 1:200 dilution), which targets the 3Myc epitope, and probed with Alexa Fluor 546-conjugated goat anti-mouse secondary antibody. Scale bars: 5 µm. (C) Immunogold labelling of 3Myc::mNG::PFR2 (positive control) and untagged parental (negative control) cell lines. Insets show a detailed view of the different structures, highlighting a high concentration of 10 nm gold particles (arrowheads) in the PFR for the mNG::3Myc::PFR2 tagged cell line, with few or no gold particles in the filaments for the untagged cell line. PFR: paraflagellar rod. (D) Quantification of the number of gold particles in the filaments (top) and PFR (bottom) for the parental, KIAP2::3Myc::mNG and 3Myc::mNG::PFR2 cell lines. Images (parental: n = 6; KIAP2::3Myc::mNG:: n = 7; 3Myc::mNG::PFR2: n = 6) were partitioned into 1 $\mu m^2$ grids in Fiji; four complete squares were achieved for each image. The number of gold particles in grids with filaments and PFR were counted and the average gold particle count in the structures plotted. The horizontal bars represent the mean of gold particles in each structure across the cell lines. P-values were calculated using the Mann-Whitney test.

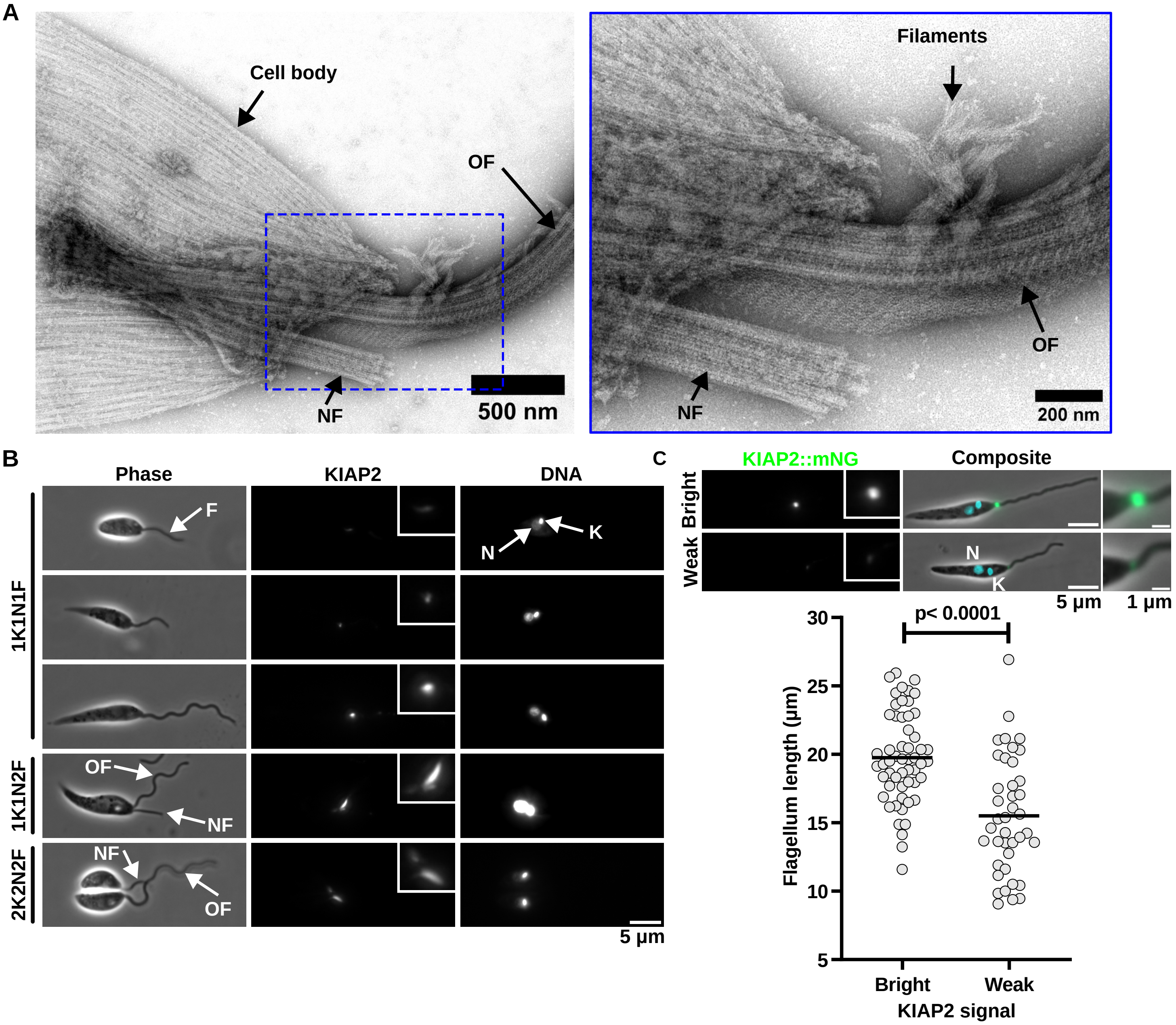


**Fig S5. Flagellar filaments and KIAP2 signal intensity at different stages of *L. mexicana* cell cycle.** (A) Negatively stained dividing cell showing the presence of the filaments on the old flagellum (OF), but not on the new flagellum (NF). (B) Fluorescence micrographs of major cell cycle stages based on the number of kinetoplasts (K), nuclei (N) and flagella (F). KIAP2 was C-terminally tagged with mNG, while the nucleus and kinetoplast were stained with Hoechst 33342. Insets show a magnified view of the KIAP2::mNG signal in the flagellum of cells in each stage. In the 2F cells, KIAP2 signal was much brighter in the old flagellum (OF) compared to the new flagellum (NF). Scale bar: 5 µm. (C) Quantification of KIAP2 signal in 1K1N1F cells (bright: n = 55; weak: n = 40). We determined the flagellum lengths by measuring the distance between the kinetoplast and the tip of the flagellum using Fiji. We calculated the p-values using the Mann-Whitney test.

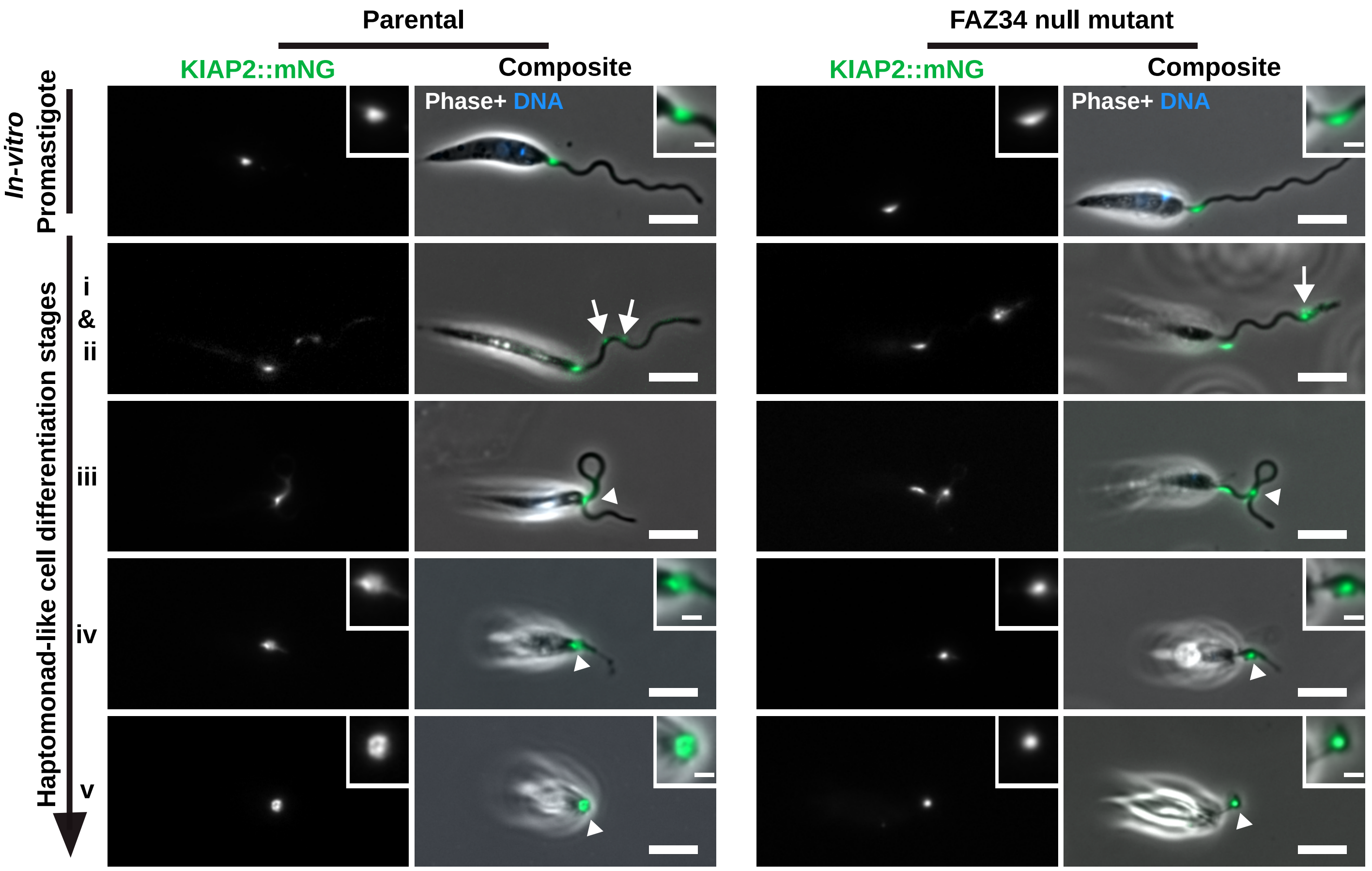


**Fig S6. Deletion of FAZ34 separated the cell body from the KIAP2 signal and the adhesion plaque during adhesion of haptomonad-like cells.** Development of mNG-tagged KIAP2 in the parental (left) and FAZ34 null mutant (right) *L. mexicana in vitro* promastigotes and haptomonad-like cells. The arrows show bright spots associated with initial foci of adhesion,while the arrowheads denote the position of the adhesion plaque in haptomonad-like cells. The composite shows the phase contrast, KIAP2::mNG (green) and DNA (blue) signals. Scale bars: 5 µm; insets: 1 µm

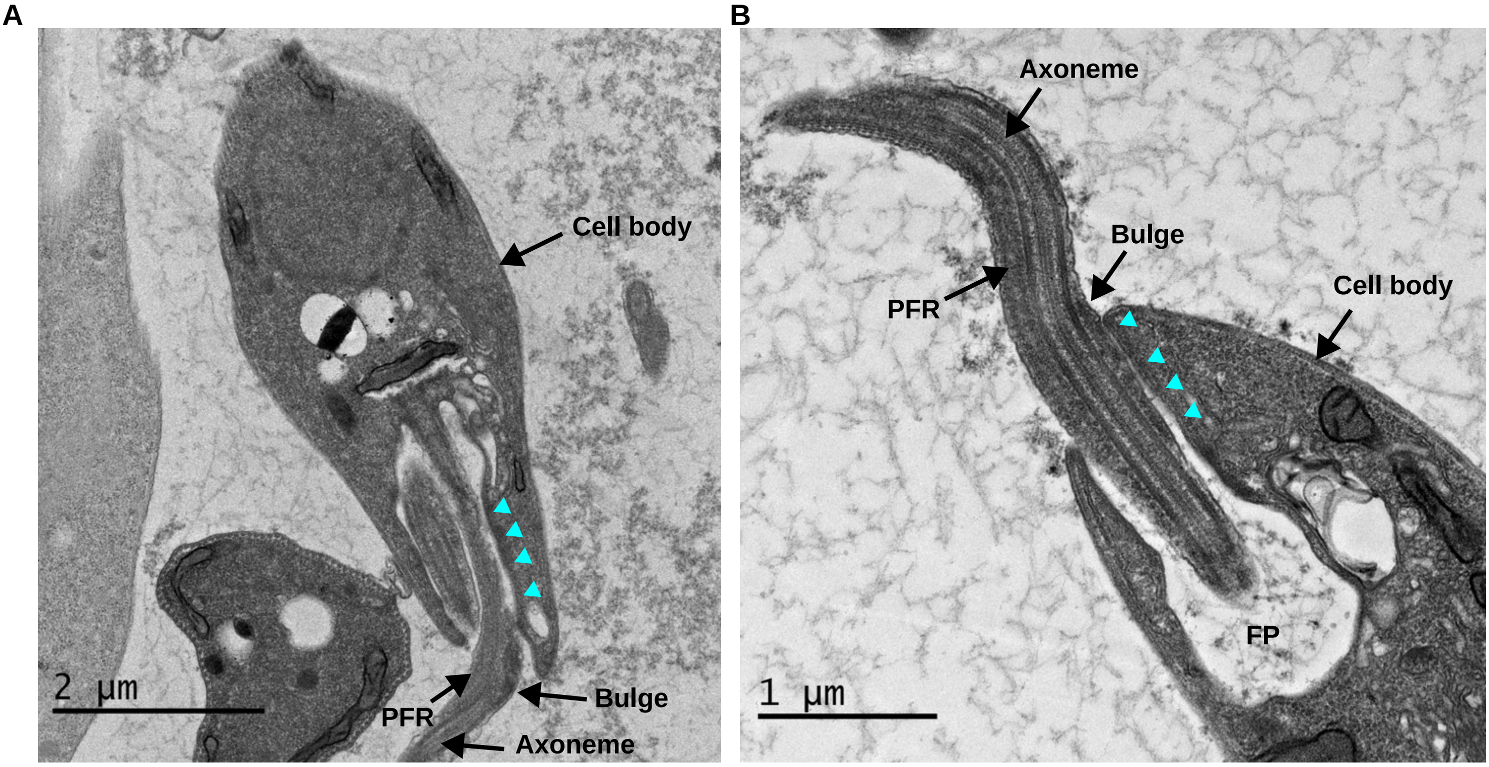


**Fig S7. Thin-section electron micrographs of flagellar filaments in *L. mexicana* promastigote.** (A-B) Transmission electron micrographs of *L. mexicana* longitudinal sections through the flagellum and flagellar pocket showing a bulge on the side associated with the flagellum attachment zone (FAZ; arrowheads), corresponding to the position of the filaments, as the flagellum emerges from the cell body. FP: Flagellar pocket
